# Mosquito Specie Composition and Pyrethroid Resistance Status in Daura, Maiadua and Sandamu LGAs, Katsina State

**DOI:** 10.64898/2026.01.09.698587

**Authors:** Abubakar Sani, Musa Yalwa Abubakar, Muhammad Musa Ahmad, Usman Salisu Batagarawa, Muhammad Auwal Abdullahi

## Abstract

**Background:** Malaria and other mosquito-borne diseases remain major public health in Nigeria. Insecticide-based interventions are vital for disease control but their effectiveness is threatened by the limited knowledge of local mosquito species and their resistance to insecticides.

**Aim:** The aim of this research is to determine the specie composition and pyrethroid resistance status of mosquitoes in Daura, Maiadua and Sandamu local governments, Katsina state.

**Materials and Method:** Mosquito larvae were collected from different locations in the study area between July to October and reared to adults in general Biology lab, federal polytechnic Daura. The adult mosquitoes were identified and subjected for bioassay. Five different WHO insecticide-impregnated papers namely Alphacypermetrin, Deltametrin, Lambdacyhalothrin and Permethrin were used for this study. The impregnated papers were tested against the emerged adult mosquitoes using WHO standard operating procedure. *Culex* and *Aedes* were morphologically identified to specie level while *Anopheles* were molecularly identified. The kdr mutation was also assessed in the *Anopheles* mosquitoes.

**Results:** All the three genera; *Anopheles*, *Culex* and *Aedes* were present in the study area with *Anopheles* being the dominant. *Culex quinquifasciatus* and *Aedes aegypti* were the dominant species of *Culex* and *Aedes* respectively, while *Anopheles gambiae s.l.* was the dominant *Anopheles* species. Bioassays revealed widespread resistance to all the tested insecticides across all genera and locations, with particularly high resistance to deltamethrin, (mortality rates 17-48%). molecular analysis showed a high frequency of the 1014 mutation in the *Anopheles* mosquito.

**Conclusion and Recommendations:** *Anopheles gambiae s. l., Culex quinquifasciatus* and *Aedes aegypti* are the dominant mosquito species in the study area. Deltametrin was less effective than the other classes of pyrethroids. These findings are important for guiding malaria control programs in Nigeria particularly in selecting effective insecticides for vector management.

## INTRODUCTION

Malaria continues to be one of the most important vector-borne parasitic diseases posing major public-health challenges of our time threatening the health of about two-third of the world’s human population in 106 countries and territories. According to [1], in 2022 there were approximately 249 million malaria cases and over 600,000 deaths globally with more than 90 % of those occurring in the African region. Each day accounts to 1000 mortality of young lives making malaria to be the 3rd leading cause of death for children under five years worldwide, after pneumonia and diarrheal disease [2]

Nigeria is estimated to have 97% of its population at risk of malaria, and malaria transmission occurs across almost the entire country; furthermore, Nigeria bears the highest malaria burden globally, accounting for ∼27% of global cases, [3,4] There are an estimated 100 million malaria cases with over 300,000 deaths per year in Nigeria, and in 2016 the country accounted for 27% of global malaria cases and 24% of malaria deaths.[5,6] In some regions of Nigeria, malaria has been reported to account for approximately 66% of outpatient visits, particularly in primary health care centers. [7]

Malaria in sub-Saharan Africa is largely transmitted by species within the *Anopheles gambiae* complex, which includes *An. gambiae s.s.*, *An. coluzzii*, and *An. arabiensisAn. arabiensis* [8] In Nigeria, a country-wide *Anopheles* vector database identified nearly 29 species, with members of the An. *gambiae* complex dominating the vector population.[9] Although *An. funestus* plays a less frequent role compared to the *gambiae* complex, it remains an important secondary vector. [10]

Prevention of vector-borne diseases is particularly challenging in the absence of widely available vaccines, making vector control and personal protection such as insect-treated nets and repellents the mainstays of prevention.[11,12] Vector control interventions, notably long-lasting insecticidal nets (LLINs) and indoor residual spraying (IRS), depend heavily on chemical insecticides,[13,14] with pyrethroids (especially permethrin and deltamethrin) dominating usage in Africa (approximately 89.9% of insecticide use by standard spray coverage).[14]

In 2023, a total of 255 million insecticide-treated nets (ITNs) - the primary vector control tool - were distributed in malaria-endemic countries, including 24 million in Nigeria.[15] Although these large-scale interventions have contributed significantly to malaria control, they have also imposed substantial selection pressure on mosquito populations, accelerating the evolution of insecticide resistance. This resistance now threatens malaria control by allowing mosquitoes to survive exposure to all major insecticide classes, particularly pyrethroids.[15,16] In line with WHO recommendations, regular monitoring of resistance profiles in highly endemic regions is essential to guide the choice of effective vector control tools.[15]

One key indicator of susceptibility is the knock-down rate (kdr), which measures how quickly mosquitoes are immobilized following insecticide exposure. The knock-down effect is an early measure of insecticide efficacy, and when KDR is prolonged or significantly reduced, it may signal emerging resistance mechanisms such as knock-down resistance (kdr) mutations or metabolic detoxification. [17,18]

Given these considerations, the knock-down rate of mosquitoes emerges as a critical entomological parameter that serves multiple functions: (1) it provides an early indicator of reduced insecticide performance, (2) it may signal the need to rotate or escalate insecticide classes, (3) it can inform local vector-control programs of the urgency for alternative interventions (e.g., larval source management, non-pyrethroid nets), and (4) it contributes to understanding residual malaria transmission in high-burden settings. In Burkina Faso, for instance, larval-source management using *Bacillus thuringiensis israelensis* (Bti) achieved a 61 % reduction in female *Anopheles* mosquitoes when breeding sites were selectively treated, and up to 70 % reduction when all breeding sites were treated. [18]

Nevertheless, despite the critical role of KDR assays, there is limited routine monitoring in many high-transmission settings particularly at sub-national levels, where vector ecology, insecticide pressure, and behavioral traits can differ substantially. A recent review of six decades of vector-control research in southern Africa identified major gaps in entomological monitoring, insecticide resistance, and the adaptability of vector-control interventions to evolving vector behaviors.[20] In Nigeria and more specifically in Katsina State published entomological data remain sparse: while *Anopheles* species composition and breeding sites have been characterized,[21] and knock down rate in *Aedes*,[22] there is little published information on knock-down rates or insecticide susceptibility in the Daura Zone. This lack of baseline resistance data limits the ability to tailor vector-control strategies to local conditions.

In light of the foregoing, this study aims to determine the specie composition and Pyrethroid resistance status in Daura, Mai;adua and Sandamu LGAs of Katsina State.

## METHODOLOGY

### Study Area

This study was carried out from July to October 2025 in three local governments; Daura, Maiadua, and Sandamu, within the Daura emirate in Katsina State. Various sites within these regions were selected for collecting mosquito larvae, particularly from stagnant water, old tires, cans, discarded clay pots, and gutters. Sampling took place between 6:00 and 8:00 am. The area experiences two main seasons: a rainy season spanning June to October, and a dry season from November to May. The majority of residents are farmers, cultivating cereals (such as millet, sorghum, and maize) during the rainy season, and vegetables (like cabbage, lettuce and pepper) during the dry season. Farmers commonly use pyrethroid-based insecticides for pest management and insecticide treated nets are distributed by the society for family health and CRC and used by the inhabitants.

**Figure.**
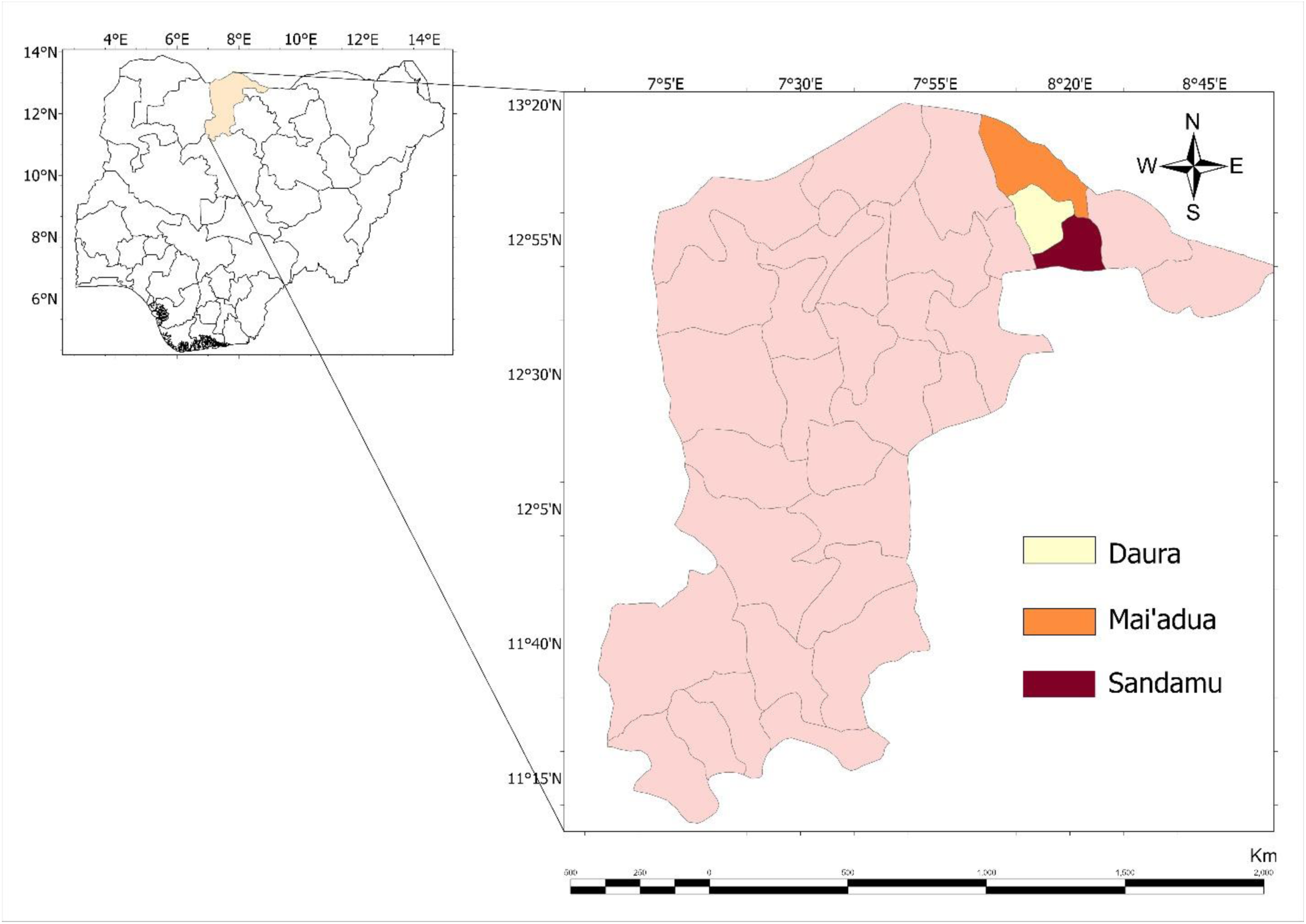
Error! No text of specified style in document.. Map of the Study Area

### Larvae Collection, Identification, and Rearing

Various methods were explored for the Mosquito larvae collection including but not limited to the dipping method with a standard 350 ml larval dipper featuring an extendable handle, taking ten dips per breeding site. The larvae were transported in clear plastic containers to the Biology Laboratory of the department of science laboratory technology, Federal Polytechnic Daura. Identification to the genus level followed the guidelines by [23]. Rearing was done in plastic jars covered with fine mesh and the larvae were fed a mixture of yeast and biscuit. Once adults emerged, they were transferred to cages and fed with sugar-soaked cotton wool. Adults aged 3-5 days were used for susceptibility testing, following guidelines.[2]

### Morphological Identification of *Anopheles Culex* and *Aedes* Species

Adult mosquitoes were identified in to genera under dissecting and compound microscope using African standard key and pictorial keys [23,24,25]. *Culex* and *Aedes* were identified to specie level using morphological keys.[25] This was done by studying the pattern of the antennae, the scales and color of the palps at the head region, the patterns of spots on the wings, thorax, terminal abdominal segments, scales of the legs, presence or absence of speckling in the hind tarsi and striations on the body using both compound and dissecting microscope following the taxonomic keys.

### Bioassay Test

Bioassays were conducted as per WHO protocols, exposing mosquitoes to insecticide-treated papers. For each insecticide, four replicates of 20-25 mosquitoes were tested. Mosquitoes were first acclimated for an hour, then placed in tubes with papers impregnated with Alpha-cypermethrin, Deltamethrin, Lambda-cyhalothrin, and Permethrin. Knockdown was recorded over an hour, after which mosquitoes were moved back to holding tubes and kept for another 24 hours, with moist cotton wool provided. Mortality was assessed after 24 hours. Controls, comprising two batches per insecticide, were subjected to untreated papers. All experiments were performed in the same laboratory, with temperature and humidity maintained at roughly 29±2°C and 67-80%.

### DNA Extraction

The morphologically identified *Anopheles gambiae* mosquitoes were carefully placed in well-labelled Eppendorf’s tube containing silica gel using forceps without damaging any part of the body. They were sent to the Africa center of excellence for neglected tropical disease and forensic biotechnology (ACENTDFB), Ahmadu Bello University Zaria, Nigeria, for molecular identification and 1014 mutation detection.

Genomic DNA extraction was performed using the Zymo Research kit as per the manufacturer’s instructions. Mosquito samples were homogenized, treated with proteinase K, and further processed with lysis buffer. After centrifugation, the supernatant was applied to a Zymo-spin column, washed, and finally eluted with DNA elution buffer.

### Molecular Identification of *Anopheles gambiae*

The extracted DNA was used for molecular identification by PCR, following [27] with slight modifications. The PCR involved 30 cycles with specific denaturation, annealing, and extension steps (denaturation at 94°C for 30sec, annealing at 50°C for 30s sec, and extension at 72°C for 30 sec). Species-specific primers (UN, GA, AR, QD) were used, resulting in distinct amplicon sizes for *An. quadriannulatus* (153bp), *An. arabiensisAn. arabiensis* (315bp), and *An. gambiae* (390bp).

### PCR Amplification for Kdr Allele

The genomic DNA extracted from mosquitoes was also used in genotyping of resistant (L1014F allele) and susceptible (L1014L) mosquitoes, using the protocol prescribed by[26] Primers sequences (Agd1, Agd2, Agd3 and Agd4) used are described in Table 1. This protocol allows the amplification of the DNA fragment coding for the voltage-dependent sodium channel in each tested mosquito. A multiplex PCR was carried out allowing the Agd1/Agd2 primers pair amplifying a 293 bp product of the *kdr* gene as an internal control. The Agd3/Agd1 primer pairs amplifies the resistance portion of the *kdr* gene as a 195pb fragment while the Agd4 /Agd2 pair amplifies a 137 bp fragment associates with the sensitive or susceptible gene. The PCR mixtures (25µl) consisted of 8.0µl of DNA template, reconstructed primers (2.0µl), master mix (12.5µl), coral load (2.0µl) and 0.5µl of nuclease free water and The PCR reaction conditions were 1 min at 94°C, 2min at 50°C and 2 min at 72°C for forty cycles with a final extension step at 72°C for 10 min.

**Table 1.**
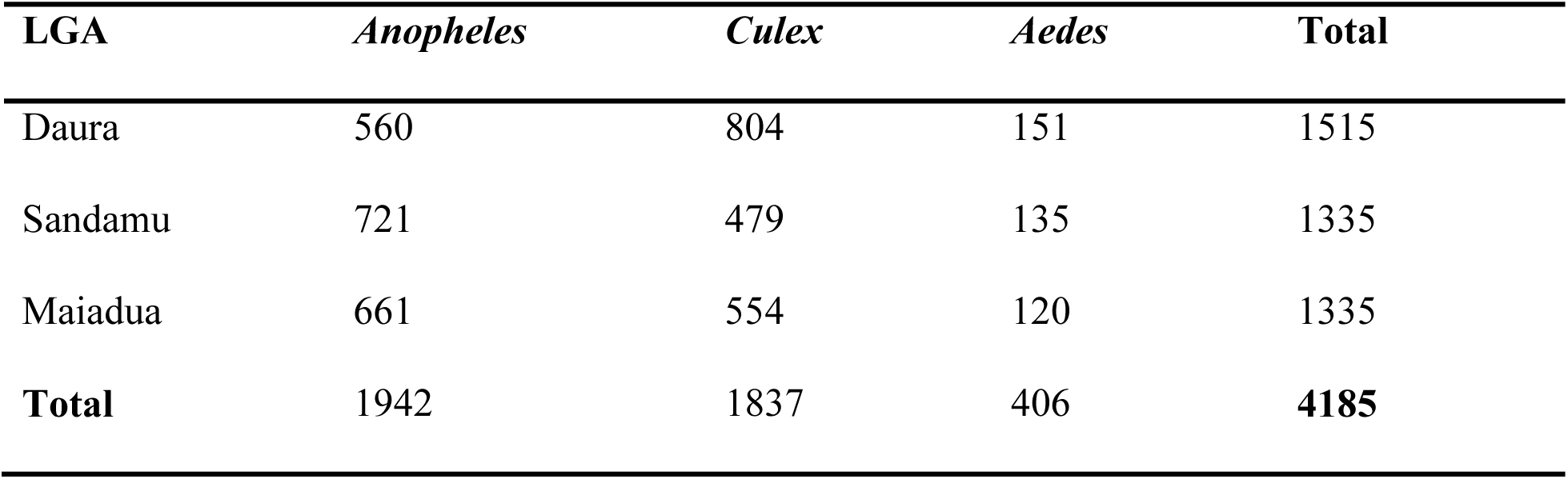
Genera Composition of Mosquitoes Collected from Three LGAs in Daura Zone.

### Agarose Gel Electrophoresis

PCR products were separated using 2% agarose gel electrophoresis. The gel was made with 70ml water and 1.08mg agarose, microwaved, and stained. Samples were loaded and run at 70V for 45 minutes, then visualized with a gel documentation system.

**Table 1.1:**
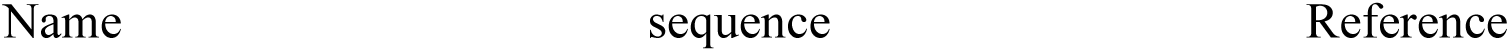
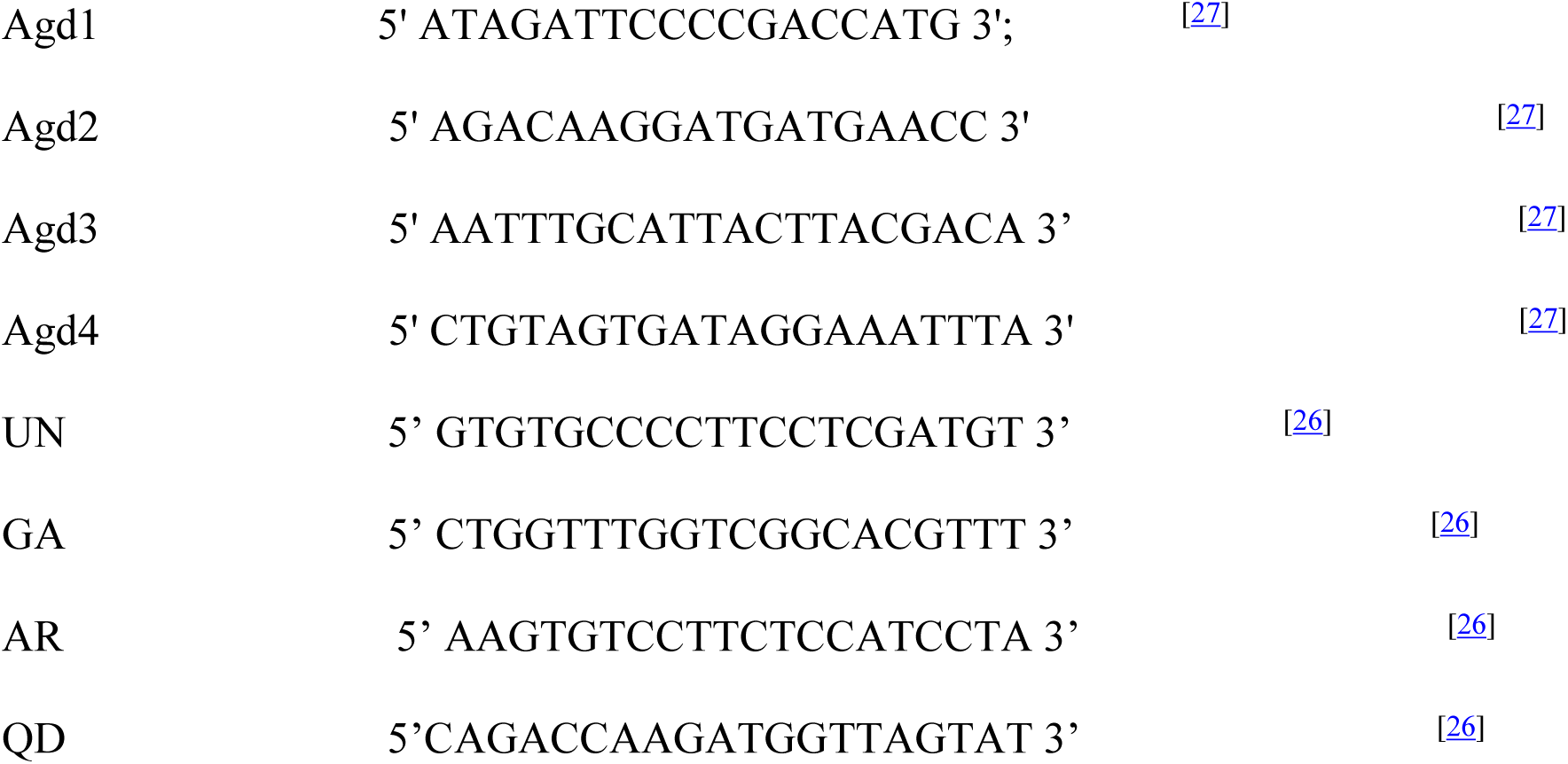
Sequences of the primers used for *Anopheles Specie* Identification and Kdr mutation genotyping.

## Results

### Mosquito genera composition

Table 1 shows the composition of mosquitoes by genera that were collected from three local governments in Daura Zone. A total of 4,185 mosquitoes belonging to three genera; *Anopheles, Culex,* and *Aedes* were recovered from the study locations. The highest number of mosquitoes were collected from Daura (n=1515). *Anopheles* was the most abundant genera in all the three local governmets.

### Mosquito species composition

Four species of mosquitoes from three genera were identified as shown in Table 2. *Anopheles gambiae* s.l. was the most prevalent species with 1,473 individuals, followed by *Culex pipiens* (n=1437) and *Aedes aegypti* (n=1,129). Other species included *Anopheles funestus* (n=69), and *Aedes albopictus* (n=77). The distribution of species was also varied by location, with *An. Gambiae s.l.* being the most abundant in Sandamu and Maiadua, meanwhile *Ae. aegypti* was the most frequent specie in Daura.

**Table 2.** Species Composition of Mosquitoes Collected from Three LGA in Daura Zone.

| Specie | Daura | Sandamu | Maiadua | Total |
| --- | --- | --- | --- | --- |
| *Anopheles gambiae* s.l. | 313 | 611 | 549 | 1473 |
| *Anopheles funestus* | 47 | 10 | 12 | 69 |
| *Culex pipiens* | 534 | 429 | 474 | 1437 |
| *Aedes aegypti* | 550 | 279 | 300 | 1129 |
| *Aedes albopictus* | 71 | 6 | 0 | 77 |
| **Total** | 1515 | 1335 | 1335 | 4185 |

### Molecular identification of *Anopheles gambiae* s.l. species

We further investigated the species composition of the *An. gambiae s.l.* complex by subsampling 60 mosquitoes (20 mosquitoes per location). Result (shown in figure 1) revealed the presence of two species within the complex; *An. gambiae s.s.* and *An. arabiensisAn. arabiensis* in all locations. Out of the total 60 mosquitoes examined across all locations, *An. gambiae s.s.* constituted 34 individuals (56.7%), while *An. arabiensisAn. arabiensis* was made up 26 individuals (43.3%) and no *An. quadriannulatus* was detected. Sandamu had the highest proportion of *An. gambiae s.s.* (14/20), whereas Maiadua had a higher number of *An. arabiensisAn. arabiensis* (11/20).

**Figure 1.**
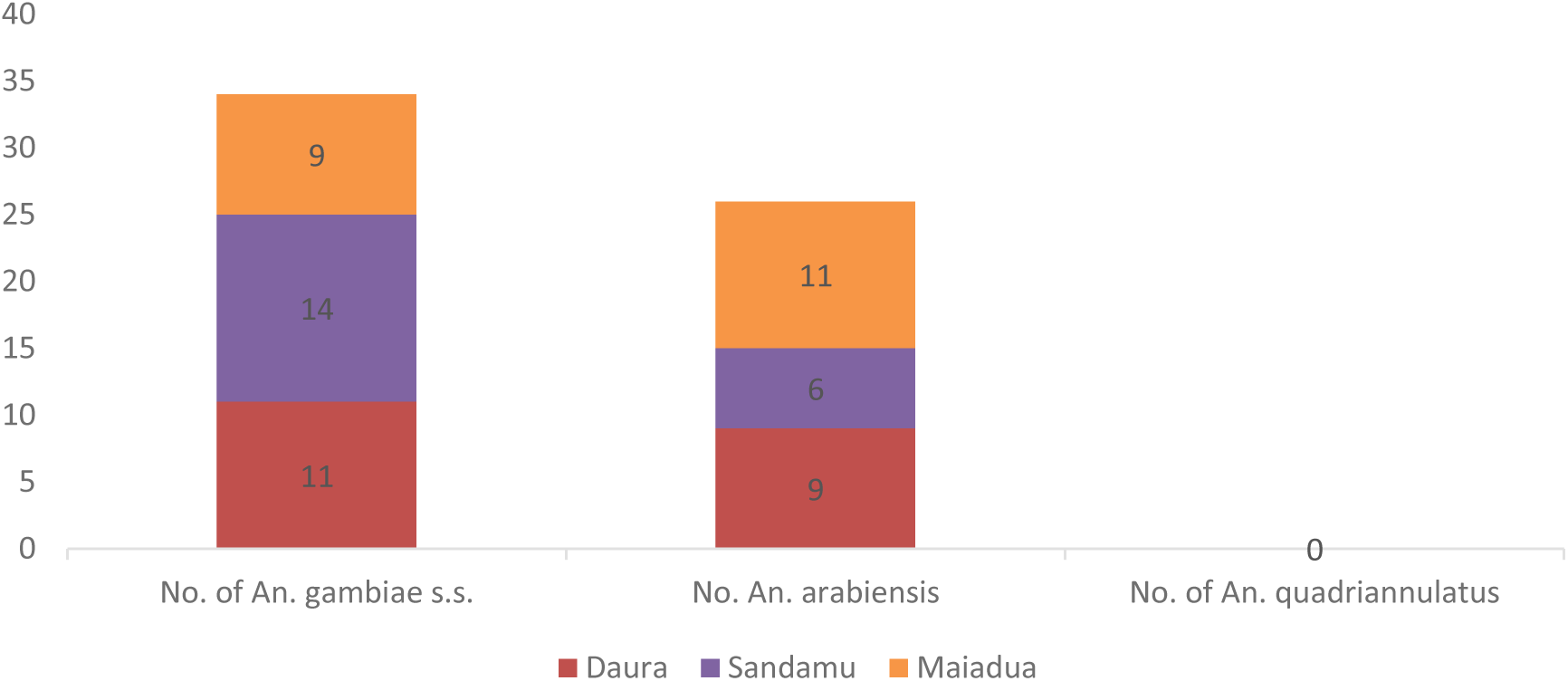
Distribution of Anopheles gambiae s.l. species

### Susceptibility Bioassay

#### *Anopheles* Mosquitoes

*Anopheles* mosquitoes showed varied resistance status to the tested insecticides as shown in Table 3. In Daura, the mosquitoes showed possible resistance to alpha-cypermethrin (97.3% mortality) and permethrin (95.0% mortality) but were deltamethrin (32.0%). Lambda-cyhalothrin showed susceptibility (98.9%). Similar patterns of resistance, possible resistance, and susceptibility were noted in Sandamu and Maiadua.

**Table 3.**
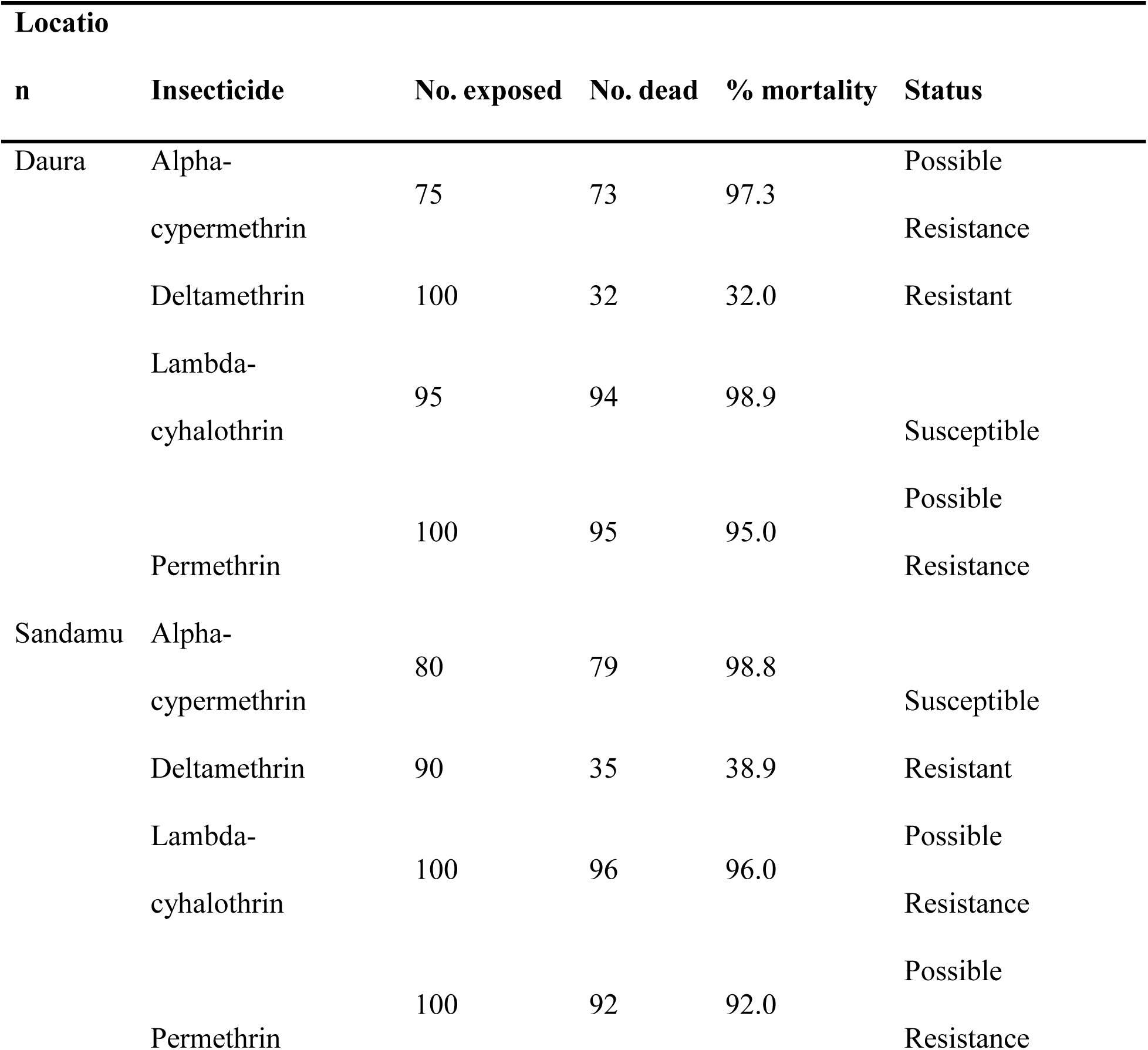
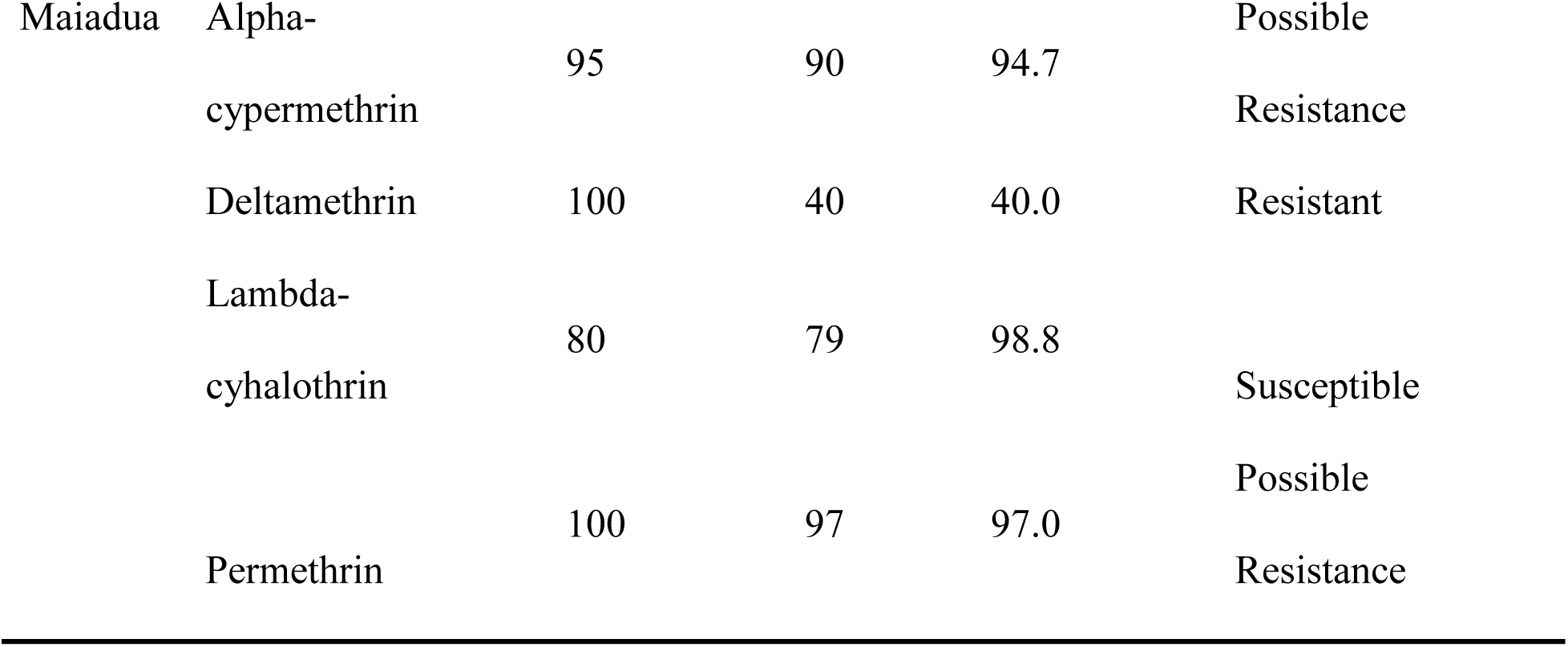
Susceptibility Status of *Anopheles* Mosquitoes exposed to five insecticides.

#### *Culex* Mosquitoes

*Culex* mosquitoes showed resistance to alpha-cypermethrin and deltamethrin across all locations (Table 4). Susceptibility was highest to lambda-cyhalothrin and permethrin, with permethrin showing 99% mortality in Daura and similar high mortalities in other LGAs. Lambda-cyhalothrin susceptibility varied from susceptible to possible resistance depending on location.

**Table 4.** Susceptibility Status of *Culex* Mosquitoes exposed to five insecticides.

| **Locatio** |  |  |  |  |  |
| --- | --- | --- | --- | --- | --- |
| **n** | **Insecticide** | **No. exposed** | **No. dead** | **% mortality** | **Status** |
| Daura | Alpha-cypermethrin | 100 | 72 | 72 | Resistant |
|  | Deltamethrin | 100 | 36 | 36 | Resistant |
|  | Lambda-cyhalothrin | 100 | 98 | 98 | Susceptible |
|  | Permethrin | 100 | 99 | 99 | Susceptible |
| Sandamu | Alpha-cypermethrin | 100 | 77 | 77 | Resistant |
|  | Deltamethrin | 100 | 43 | 43 | Resistant |
|  | Lambda- |  |  |  | Possible |
|  | cyhalothrin | 100 | 97 | 97 | Resistance |
|  | Permethrin | 100 | 98 | 98 | Susceptible |
| Maiadua | Alpha- |  |  |  |  |
|  | cypermethrin | 100 | 81 | 81 | Resistant |
|  | Deltamethrin | 100 | 48 | 48 | Resistant |
|  | Lambda- |  |  |  |  |
|  | cyhalothrin | 100 | 100 | 100 | Susceptible |
|  | Permethrin | 100 | 98 | 98 | Susceptible |

#### *Aedes* Mosquitoes

As shown in Table 5, *Aedes* mosquitoes showed susceptibility to alpha-cypermethrin in Daura (99%) and permethrin in Sandamu and Maiadua (100% and 99%, respectively). Resistance was was observed to deltamethrin, and lambda-cyhalothrin across all LGAs, with mortality rates generally below 90%. Possible resistance was recorded for permethrin in Daura and Maiadua.

**Table 5.**
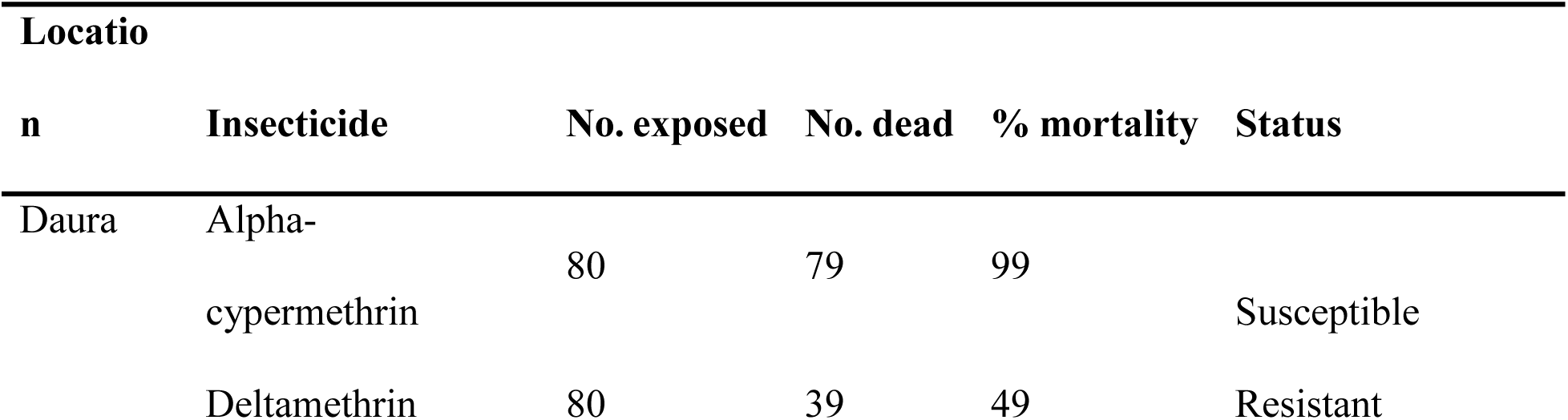
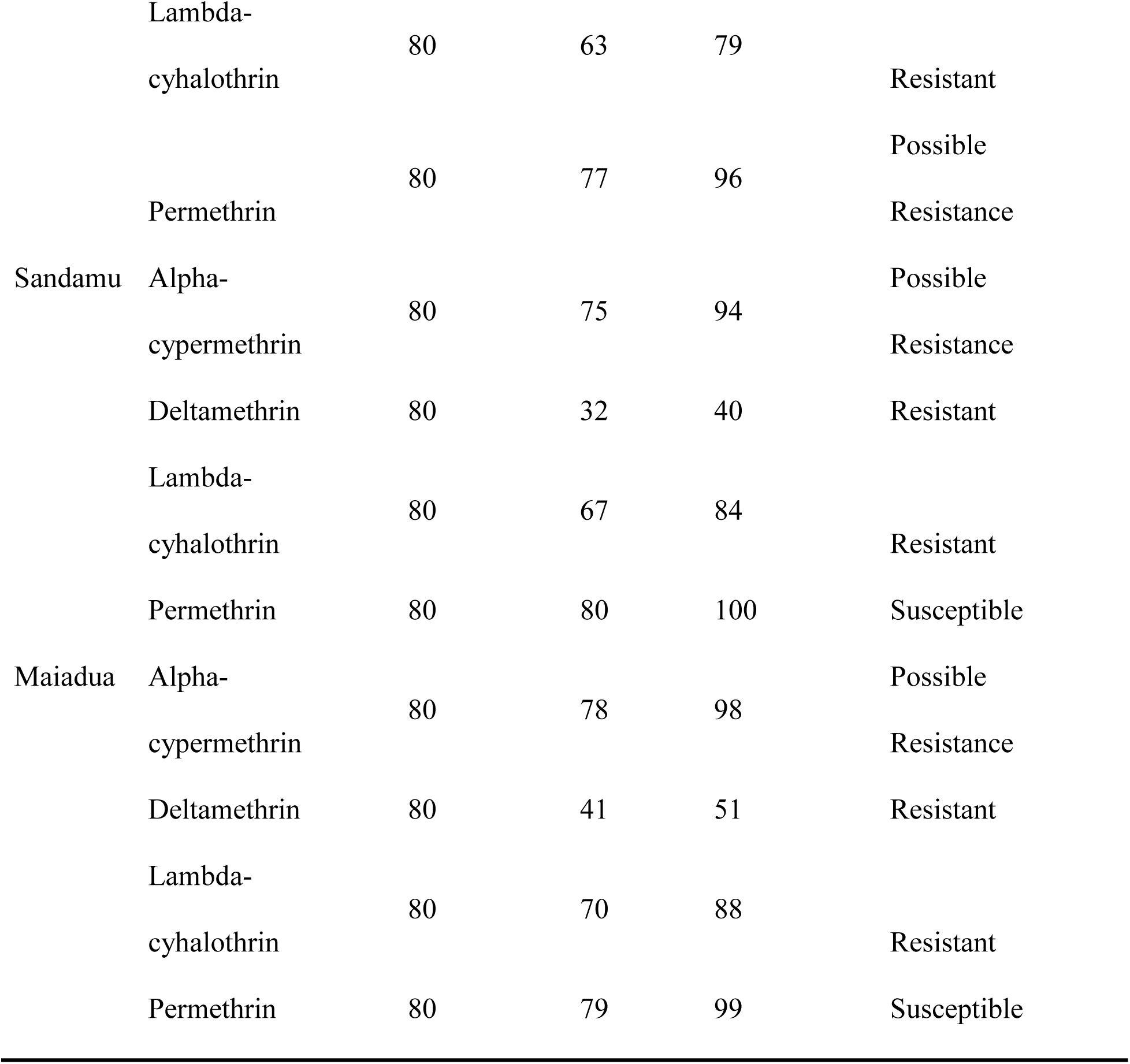
Susceptibility Status of *Aedes* Mosquitoes exposed to five insecticides.

### Frequency of kdr 1014F allele

The frequency of the kdr 1014F allele in *Anopheles* mosquitoes is presented in Table 6. There were varying frequencies for the kdr allele across the three LGAs, where the frequency ranged from 55% in Maiadua to 77.5% in Daura.

**Table 6.**
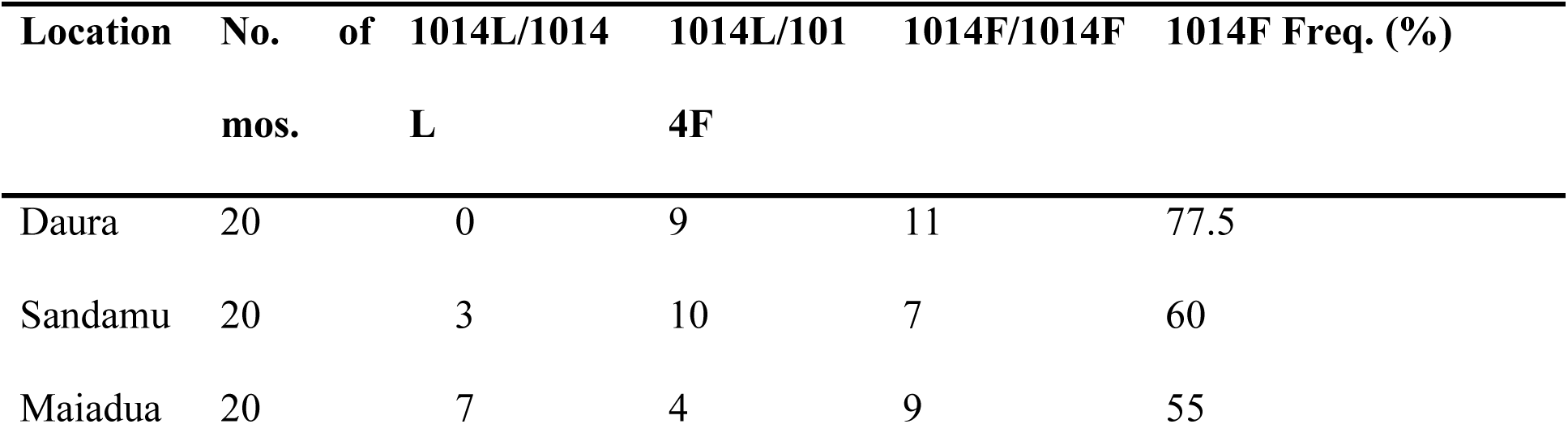
Frequency of the kdr 1014F allele in *Anopheles* mosquitoes collected from different locations in Daura Zone.

| Location | No. of mos. | 1014L/1014L | 1014L/1014F | 1014F/1014F | 1014F Freq. (%) |
| --- | --- | --- | --- | --- | --- |
| Daura | 20 | 0 | 9 | 11 | 77.5 |
| Sandamu | 20 | 3 | 10 | 7 | 60 |
| Maiadua | 20 | 7 | 4 | 9 | 55 |

## DISCUSSION

The global effort to control and eliminate malaria and other mosquito-borne diseases is increasingly challenged by the rapid development of insecticide resistance in mosquito vectors. The extensive and prolonged use of pyrethroids (accounting for about 89.9% of interventions) has fueled the emergence and spread of resistance, which undermines the effectiveness of long-lasting insecticidal nets (LLINs) and indoor residual spraying (IRS), key tools for preventing mosquito-borne diseases.[30] This issue is especially acute in high-burden areas like sub-Saharan Africa, particularly West Africa, where it weakens the impact of ITNs and IRS and contributes to roughly 96% of global malaria deaths.[1] Regular surveillance of mosquito species of health importance and insecticide resistance patterns is therefore vital to guide decisions on insecticide selection, rotation, or combination strategies,[1] support evidence-based policy development and drive innovation in next-generation vector control tools.[3] In this study, we conducted a comprehensive assessment of mosquito genera and species composition, pyrethroid resistance status and the frequency of the kdr 1014F mutation in mosquito populations from three selected LGAs in Daura Zone (Sandamu, Maiadua, and Daura).

*Anopheles* was the most abundant genus overall across the three LGAs, then *Culex*, and finally *Aedes*. However, there were notable variations within the LGAs where *Aedes* and *Culex* dominate the mosquito population in Daura LGA, which could be a result of its semi-urban environment that favors peridomestic breeding for *Aedes* and *Culex*. This prevalence could increase the risk for arboviral diseases like dengue and Zika in Daura. Further species analysis showed that *Anopheles gambiae* s.l. is the most abundant species overall across the three LGAs, showing its role as the primary malaria vector in these locations. This composition aligns with patterns observed by [31] in parts of Katsina state, where *An. gambiae s.l.* predominates due to its adaptation to temporary breeding sites. *Anopheles funestu*s was also uncovered from all the study locations, however at a significantly lower number than *Anopheles gambiae* s.l, agreeing with studies of [32] who reported that *Anopheles gambiae s. l*. was the dominant species in Gidan Yero village, Sokoto. At the same time, *Aedes aegypti* dominates the *Aedes* species across the regions. This observation is in line with previous studies conducted in other northern regions in Nigeria[22,33,34] which consistently reported *Aedes aegypti* as the predominant species in various urban and peri-urban settings. It can also be observed from the results that *Culex pipiens* is the dominant *Culex* species across the study area. This specie, a known arboviral and filarial vector, is widely spread across sub-Saharan Africa and Asian countries as reported by.[35,36,37] *Culex. pipiens* has also been documented as the most common *Culex* species in several East, West, and Central African countries, including Kenya [38], Tanzania,[39,40] Senegal [41], Benin,[42] and Zambia. [25]

Molecular identification of *An. gambiae s.l.* revealed the presence of *An. gambiae s.s.* and *An. arabiensisAn. arabiensis*, with no detection of *An. quadriannulatus*. This sibling species distribution in our study is in line with previous reports from Nigeria, where *An. gambiae s.s.* and *An. arabiensisAn. arabiensis* coexist in sympatry, often with *An. gambiae s.s.* consistently emerging as the predominant species across different ecological zones.[22,43,44] The higher proportion of *An. gambiae s.s.* in Sandamu compared to Maiadua may indicate micro ecological differences, as *An. arabiensis* is better adapted to the drier conditions prevalent in Maiadua. This composition has implications for malaria transmission dynamics, as *An. gambiae s.s.* exhibits higher anthropophily (preference for human blood feeding) and endophily (indoors feeding), potentially increasing vectorial capacity in areas like Sandamu.[45] In comparison, studies in southern Nigeria have reported a shift toward *An. coluzzii* dominance in urban settings,[46] suggesting that the Daura Zone’s profile reflects a more rural, Sahelian influence. These findings emphasize the importance of molecular tools in resolving cryptic species complexes, as morphological identification alone may underestimate transmission risks.

Insecticide susceptibility bioassays showed widespread resistance to all the tested insecticides across all genera in all locations. The observed resistance was more profound to deltamethrin, with mortality rates ranging from 17-48%. *Anopheles* populations exhibited resistance to deltamethrin, with possible resistance to permethrin, alpha-cypermethrin, and lambda-cyhalothrin in most sites. This is consistent with previous pyrethroid resistance documented in *Anopheles* mosquitoes across Nigeria. For instance, similar high-level resistance to deltamethrin has been reported in southern Nigeria, attributed to target-site mutations and metabolic detoxification.[17] In Katsina State specifically, recent investigations in Musawa LGA have confirmed resistance in *Anopheles* to permethrin, although with variable susceptibility to organophosphates, which was not tested in our study.[47] *Culex* and *Aedes* genera showed similar resistance status, with uniform resistance to alpha-cypermethrin, and deltamethrin, while showing higher susceptibility to permethrin and lambda-cyhalothrin. Comparable resistance has been noted in parts of Nigeria, including Katsina State.[22,48,49] The consistent resistance across genera as observed in our study indicates shared environmental pressures, such as indiscriminate insecticide use in farming, which could undermine Nigeria’s National Malaria Elimination Program as well as the control of other mosquito-borne diseases.

Molecular detection of kdr mutation revealed a high presence of the 1014 mutation in the *Anopheles* mosquito from the study area. This presence of high resistance gene can be attributed to the long term usage of insecticide treated nets (ITNNs) that are been distributed every year in the areas and the fact that inhabitants of the area mostly engaged in farming activities, and there is rampant and continuous usage of agricultural chemicals in the farms. These chemicals are sometimes washed deposited in the surrounding water bodies which serve as breeding ground for the mosquitoes. The developing larvae might get exposed to the chemical residues thereby enhancing development of resistant. Therefore, the combination of consistent ITN use and extensive agricultural pesticide application in the field may synergistically contribute to the observed resistance patterns.

